# A steroid hormone regulates growth in response to oxygen availability

**DOI:** 10.1101/2021.06.25.449962

**Authors:** George P. Kapali, Viviane Callier, Hailey Broeker, Parth Tank, Samuel J.L. Gascoigne, Jon F Harrison, Alexander W. Shingleton

**Affiliations:** Department of Biological Sciences, University of Illinois Chicago, 840 West Taylor Street Chicago, IL 60607; School of Life Sciences, Arizona State University, Tempe, AZ 85287-4501; Department of Biology, Lake Forest College, 555 North Sheridan Road, Lake Forest, Illinois 60045

## Abstract

In almost all animals, physiologically low oxygen (hypoxia) during development slows growth and reduces adult body size^1–3^. The developmental mechanisms that determine growth under hypoxic conditions are, however, poorly understood. One hypothesis is that the effect of hypoxia on growth and final body size is a non-adaptive consequence of the cell-autonomous effects of hypoxia on cellular metabolism. Alternatively, the effect may be an adaptive coordinated response mediated through systemic physiological mechanisms. Here we show that the growth and body size response to moderate hypoxia (10% O^2^) in *Drosophila melanogaster* is systemically regulated via the steroid hormone ecdysone, acting partially through the insulin-binding protein Imp-L2. Ecdysone is necessary to reduce growth in response to hypoxia: hypoxic growth suppression is ameliorated when ecdysone synthesis is inhibited. This hypoxia-suppression of growth is mediated by the insulin/IGF-signaling (IIS) pathway. Hypoxia reduces systemic IIS activity and the hypoxic growth-response is eliminated in larvae with suppressed IIS. Further, loss of *Imp-L2*, an ecdysone-response gene that suppresses systemic IIS, significantly reduces the negative effect of hypoxia on final body size. Collectively, these data indicate that growth suppression in hypoxic *Drosophila* larvae is accomplished by systemic endocrine mechanisms rather than direct suppression of tissue aerobic metabolism.

---

The hypothesis that the effect of hypoxia on growth and final body size is regulated cell-autonomously rests on the general phenomenon whereby, in respiring cells, oxygen-limitation reduces the ability of the cell to generate ATP, as cells switch from aerobic to anaerobic respiration. This in turn slows the biological processes that rely on ATP, which include cell growth and proliferation. To determine what level of hypoxia induces anaerobic metabolism, we looked at the accumulation of lactate, the primary anaerobic end-product in *Drosophila*^4^, in *Drosophila* larvae transferred from 21% oxygen to 3%, 5% and 10% oxygen in the third larval instar. We found that, while transfer to 5% and 3% oxygen resulted in strong and prolonged accumulation of lactate, no such accumulation was observed in larvae transferred to 10% (Fig 1 A). Nevertheless, larvae transferred to 10% oxygen at ecdysis to the third larval instar show a marked reduction in overall body size of 31% (Fig 1 B, B’s), suggesting that growth suppression in 10% oxygen occurs without limits on aerobic ATP production. Different organs show a different size response to hypoxia, however. Rearing at 10% O_2_ has the same effect on the size of the wing, leg, thorax and mouthparts of the fly, while the size of the male genitalia is largely unaffected (Supp Fig 1). Intriguingly, the pattern of organ-size reduction in response to hypoxia is statistically indistinguishable from the pattern in response to low nutrition, but very different from the pattern in response to an increase in rearing temperature, which also reduces body size (Supp Fig 1). Collectively, these data suggest that the effect of 10% oxygen on growth and final body size is mediated through a systemic rather than cell-autonomous mechanism, and that this mechanism overlaps with the physiological mechanisms that slow growth and reduce final body size in response to low nutrition.

**Figure 1:**
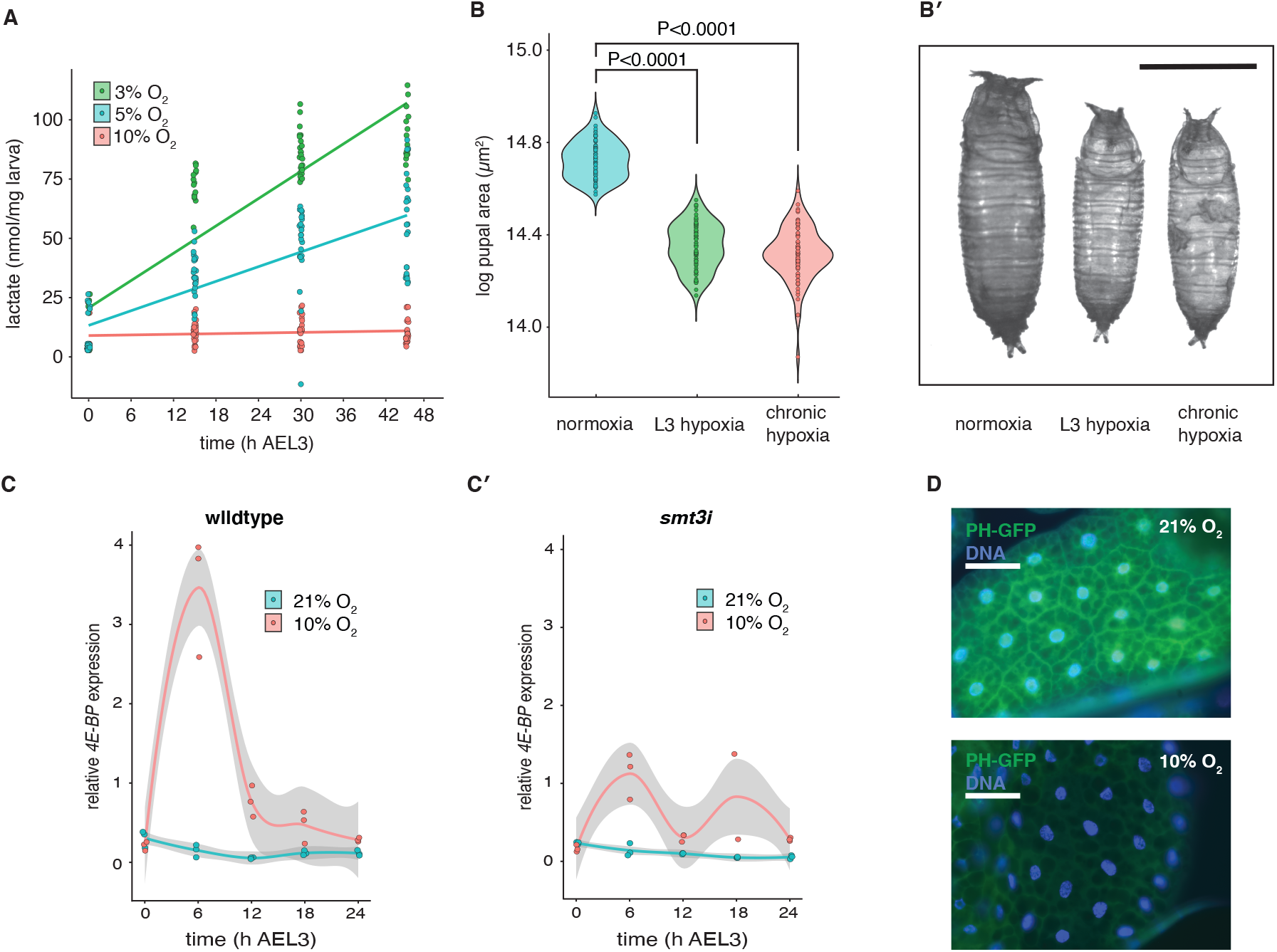
Moderate hypoxia reduces body size by suppressing IIS activity but without limits on ATP production. (A and Aʹ) Lactate accumulation in third instar larvae transferred to different oxygen levels. Transfer to 3% and 5% O_2_ resulted in significant increase in lactate (OLS regression, *P* < 0.0001 for both). Transfer to 10% O_2_ had no effect on lactate levels (OL regression, *P* = 0.1). (B) Final body size (pupal case area) is reduced relative to normoxia when flies are reared in hypoxia throughout development (chronic hypoxia) or when reared in hypoxia only during the third larval instar (L3 hypoxia). Body size between treatments was compared using pooled *t*-tests. (B’
s) Black bar is 1mm. (C) Hypoxia increases expression of *4E-BP* in the first 24h after ecdysis to the third larval instar (AEL3) relative to normoxic controls. (C’) This increase in expression is alleviated in *smt3i* larvae, which have reduced ecdysone levels. (D) TGPH membrane localization in hypoxic and normoxic conditions. TGPH localizes to the cell membrane when IIS-pathway activity is high. White bar represents 100µm. Grey bands are 95% confidence limits around the mean.

In *Drosophila* and almost all other animals, the growth response to nutrition is regulated systemically by circulating insulin-like peptides (called dILPs in *Drosophila)*, with different tissues showing different sensitivities to the effect of insulin-like peptides on growth^5^. The effect of dILPs on growth is mediated by the highly conserved insulin/insulin-like-growth-factor signaling (IIS) pathway^6,7^. In *Drososphila*, dILPs bind to the sole insulin receptor (dInR) of proliferating cells and initiate a downstream kinase cascade that includes phosphatidylinositol 3-kinase (PI3K) and serine-threonine protein kinase (Akt)^6,8–14^. Under well-fed conditions, activation of IIS-pathway increases Akt activity, which promotes growth in part by inhibiting the activity of the Forkhead Box class O transcription factor FOXO. This in turn prevents FOXO from upregulating the expression of a number of growth inhibitors, including the translation inhibitor 4E-BP^15–18^. To test the hypothesis that hypoxia reduces growth by suppressing IIS, we first looked at the effect of hypoxia on body size in flies carrying a mutation of *InR* that causes constitutive suppression of IIS activity. Hypoxia had no effect on their final body size (Supp Fig 2), supporting the hypothesis that changes in IIS activity are necessary for the growth response to hypoxia. We then measured the mRNA levels of *4E-BP* under normoxic (21% O_2_) and hypoxic (10% O_2_) conditions (Fig. 1 C). *4E-BP* expression is high when IIS activity is suppressed, and we observed an increased *4E-BP* expression when larvae were reared in hypoxic conditions. We also looked at the effect of hypoxia on Pi3K activity using the TGPH reporter construct, which localizes to the cell membrane when Pi3K and IIS-activity is high ^19^. We found that Pi3K activity was low at 10% O_2_ (Fig 1 D). Collectively, these data show that the IIS-pathway is suppressed under hypoxic conditions.

Previous studies have shown that hypoxia elevates levels of circulating ecdysone^20^. In holometabolous insects like *Drosophila*, ecdysone is a key developmental regulator that controls molting and the timing of metamorphosis^2,20–24^. However, it is also known that elevated basal levels of ecdysone suppress growth, in part by suppressing the IIS-pathway ^25–28^. A compelling hypothesis is that the effect of hypoxia on growth is mediated through the inhibitory effect of ecdysone on growth and final body size. To test this hypothesis, we genetically ablated the prothoracic gland (PG), the organ which synthesizes ecdysone, and measured larval growth under normoxic (21% O_2_) and hypoxic conditions (10% O_2_) (Supp Fig 3). While control flies larvae reduced growth under hypoxic compared to normoxic conditions this suppression of growth was alleviated in larvae lacking a PG (Supp Fig 3). This suggests that growth suppression by hypoxia requires a functional PG in *Drosophila*. To confirm our hypothesis that ecdysone synthesis itself is required to suppress growth under hypoxic conditions, we adopted a more targeted approach by genetically suppressing ecdysone synthesis. We achieved this by specifically silencing the Drosophila Small Ubiquitin Modifier (SUMO) 2/3 homologue (smt3) in the PG through RNAi knockdown, which prevents the production of high levels of ecdysone in the third larval instar (Supp Fig 4)^29,30^. Suppression of ecdysone synthesis in the *smt3i* larvae alleviated the growth reduction caused by hypoxia (Fig 2 A’), indicating that ecdysone synthesis in the PG is necessary for hypoxic growth reduction. We next tested whether inhibiting the elevation of ecdysone under hypoxic conditions also inhibits the suppressive effects of hypoxia on IIS activity (Fig 1 C’s). We observed that in *smt3i* larvae with suppressed ecdysone synthesis, hypoxia no longer increased the expression of *4E-BP*. Together these data indicate that that ecdysone synthesis is necessary to suppress activity of the IIS-pathway under hypoxia.

**Figure 2:**
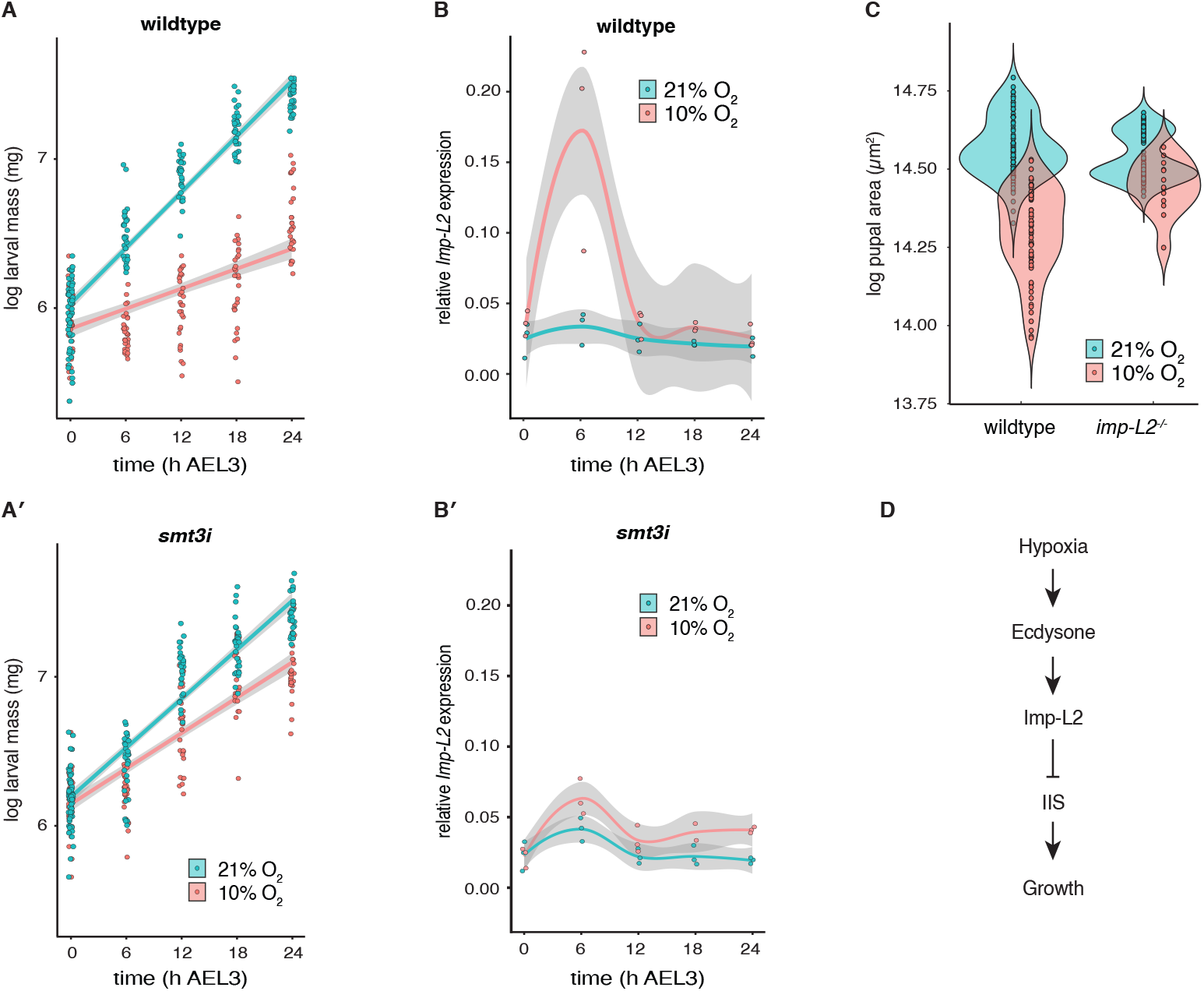
Suppression of ecdysone alleviates hypoxic growth reduction via *Imp-L2*. (A) Growth rate of wildtype third instar larvae is substantially reduced in hypoxia relative to normoxic controls. (A’) The effect of hypoxia on growth rate is significantly reduced in *smt3i* flies (GLM, *P*_*time*genotype*oxygen*_ <0.0001). (B) Expression of *Imp-L2* is elevated 6h AEL3 in wildtype hypoxic larvae. (B’) Expression of *Imp-L2* is reduced in *smt3i* hypoxic larvae. (C) Loss of Imp-L2 (*Imp-L2*^*-/-*^), almost eliminates the effect of hypoxia on final body size (*t*-test, *P*_*genotype*oxygen*_ <0.0001). (C) Model of the physiological mechanism by which hypoxia regulates growth and final body size in *Drosophila*. Grey bands are 95% confidence limits around the mean.

We next asked how ecdysone suppresses IIS and growth under hypoxic conditions. An earlier study had shown that elevated ecdysone suppresses IIS and growth in response to nutritional stress, by stimulating the release of Imp-L2 from the fat body^31^. Imp-L2 is an insulin-binding protein that antagonizes IIS systemically^32^. We therefore tested the hypothesis that Imp-L2 mediates the effect of ecdysone on growth and final body size in hypoxia. First, we confirmed that *Imp-L2* expression was transiently elevated under hypoxia (Fig 2 B), and that this elevation was suppressed in the absence of ecdysone (Fig 2 B’
s). We then tested whether loss of Imp-L2 alleviated the negative effects of hypoxia on final body size by utilizing an *Imp-L2* null mutant fly line (*Imp-L2*^*Def42*^) and found that it did (Fig 2 C). These data support a link between ecdysone, growth and final body size at low oxygen levels, where hypoxia increases ecdysone synthesis, which up-regulates expression of *Imp-L2*, which in turn suppresses growth and reduces final body and organ sizes by suppressing IIS (Fig 2 C). This is supported by the observation that ubiquitous over-expression of *dInR* alleviates growth suppression at 3.5% O_2_ in second instar larvae ^33^.

A previous study has implicated HIF-signaling in the fat body as a regulator of growth in response to low oxygen in *Drosophila*^34^. HIF-1α is a transcription factor that is activated by hypoxia^35–38^. Inducing hypoxia in the fat body alone, by limiting tracheal growth, represses dILP2 secretion from the brain, lowers systemic IIS and reduces growth, and this effect is phenocopied in normoxia by activating HIF-1α in the fat body^34^. Further, knock-down of HIF-1α in the fat body at low oxygen levels alleviates the effects of hypoxia both on dILP2 secretion and on growth^34^. What role elevated ecdysone synthesis plays in this mechanism is unclear. One hypothesis is that ecdysone suppresses growth by activating downstream components of HIF-signaling in the fat body. A key target of HIF-signaling is lactate dehydrogenase (*Ldh/Imp-L3*), an essential enzyme in anaerobic respiration that is also an ecdysone response gene^38–40^. Thus there is at least one example of crosstalk between HIF-signaling and ecdysone-signaling. An alternative hypothesis is that ecdysone and HIF-signaling act through two independent – and possibly redundant – mechanisms in the fat body to suppress IIS and slow growth in response to low oxygen. Support for this hypothesis comes from the observation that neither loss of ecdysone nor HIF-1α in the fat body appears to completely return growth in hypoxia to normoxia levels.

Our data show that ecdysone is necessary to suppress growth in hypoxia, much like for low nutrition ^31^. However, we do not have a clear understanding of how ecdysone synthesis is induced under hypoxia. During the third larval instar, timed pulses of ecdysone –interspersed with lower basal levels of ecdysone – drive key developmental events that lead to metamorphosis^41,42^. The initiation of these pulses coincides with attainment of a critical weight ^43^, at which point ecdysone synthesis becomes self-regulated by positive and negative feedback loops ^44^. Previous studies have shown that low nutrition both increases basal levels of ecdysone to slow growth but delays attainment of critical weight to delay development ^31^. The same appears to be happening in hypoxia: We measured the expression of *E74B*, an ecdysone-response gene that increases in expression in response to the ecdysone pulse that drives salivary gland histolysis ∼20h after ecdysis to the third instar. We found that low oxygen suppressed the expression of *E74B* in the first 24 hours of the third larval instar (Supp Fig 4 A), consistent with the observation that hypoxia delays attainment of critical weight and the timing of metamorphosis^45^. Nevertheless, the delay in the pulses of ecdysone cannot account for slow growth in hypoxia since loss of ecdysone synthesis increases growth rate at low oxygen. Thus it appears to be the elevated basal levels of ecdysone that slow growth at low oxygen levels ^45^.

Collectively, these data support a model where hypoxia suppresses IIS and growth in *Drosophila* via the hormone ecdysone (Fig 2 D). Loss of ecdysone alleviates the suppressive effects of hypoxia on growth rate, which indicates that larvae reared at low oxygen levels grow at a rate slower than that which can be supported metabolically. Thus, the effect of low oxygen level on growth appears to be a coordinated response regulated via a systemic mechanism rather than due solely to the cell-autonomous effects of hypoxia on cell respiration. Interestingly this systemic mechanism utilized under low oxygen shares key characteristics to the mechanism that slows growth under low nutrition; that is, ecdysone mediated growth suppression via the IIS. The fact that both low oxygen and low nutrition appear to share mechanisms to suppress growth and body size suggests that both form part of a general response to environmental stress. Consequently, animals such as *Drosophila* may have evolved generalized systemic growth-regulatory mechanisms that can respond to a multitude of stresses imposed by their ever-changing environments.

## Methods

### Drosophila Stocks

Flies were reared on standard *Drosophila* yeast cornmeal molasses medium (ref). The following fly strains were used for this study: the isogenic *RNAi* control (*60,000*) was obtained from the Vienna Drosophila RNAi Center, *phm-GAL4* (a gift from Michael O’Connor), *UAS-smt3i*^*20*^*;UAS-Hr46i* and *tub-Gal80*^*ts*^*;phm-Gal4* (gifts from Rosa barrio), *UAS-Grim* (a gift from Christen Mirth), *Imp-L2*^*def42*^ (a gift from Seogang Hyun), *P{tGPH}* (BDSC 8164), *InR*^*E19*^ (BDSC 9646), *InR*^*GC25*^ (BDSC 9554).

### Hypoxia Treatment

Unless otherwise stated, all flies were reared on standard cornmeal-molasses medium at 25°C, 21% atmospheric O_2_ level until ecdysis to the third instar. Larvae were then either maintained at 21% O_2_ (normoxic flies) or moved to 10% O_2_ (hypoxic flies).

### Lactate Assay

The lactate assay was conducted by measuring NADH produced from lactate in presence of lactate dehydrogenase (LDH) and NAD^+ 46^ NADH was measured using a Farrand Optical Components & Instruments Standard Curve System Filter Fluorometer (Valhalla, NY, USA) set at 360nm excitation wavelength and 460 nm emission wavelength.

#### Buffer preparation

To make up 100 ml of 2x Hydrazine-Tris buffer, 20 ml of a 1.0 M stock solution of Tris-Base was added to 20 ml of ddH_2_O. 10 ml of 20 M liquid hydrazine and 2.25 ml of a 125 mM stock solution of EDTA were then added to the solution, which was then bought to a pH of 10.0. ddH_2_O was added to a final volume of 100.0 ml. Finally 5 mM of NAD^+^ was added to the buffer solution right before the assay.

#### Sample preparation

50uL of HClO_4_ (17.5%) per mg of larva was added to each Eppendorf tube containing larval sample to stop metabolism and ground with a pestle. The protein was centrifuged down (1 min at 14,000g). To neutralize the acidified extract, 1.0 ml of the supernatant was added to 0.225 ml KOH + MOPS solution ([KOH]=2.0 N and [MOPS] = 0.3M)

#### Assay procedure

Each cuvette contained 20 uL of sample, 100uL of Hydrazine-Tris buffer with 5 mM NAD^+^, and 3% LDH in 90 uL ddH_2_O. Cuvettes were incubated at room temperature for 30 minutes, and then read in the fluorometer. The amount of lactate in the sample was calculated from the linear relationship between the NADH produced and lactate levels in known standard solutions.

### Nutritional, Thermal and Oxic Plasticity

We measured the effect of developmental nutrition, temperature and oxygen level on the size of five traits in wild type (*OreR*) male *Drosophila*: wing area, leg length, maxillary palp area, and the area of the posterior lobe of the genital arch. Data for the effect of nutrition and temperature on body size was taken from a previously published paper ^47^. To measure the effect of oxygen level on trait size, flies were reared throughout development at 5%, 10% or 21% O_2_. To determine the multivariate allometric coefficient under each condition, we calculated the loadings of the first eigenvector of the variance– covariance matrix for trait size, as trait size varied in response to each environmental factor. Multiplying the loadings by √*n* (where *n* is the number of traits) gives the allometric coefficient for each trait against a measure of overall body size, and is a measure of relative trait plasticity.

### Constitutive Suppression of IIS

*InR*^*E19*^*/InR*^*GC25*^ have a temperature-sensitive suppression of InR activity, such that at 17°C, InR activity is normal and at 24°C InR activity is suppressed ^48^. *InR*^*E19*^*/InR*^*GC25*^ (experimental) and *InR*^*E19*^*/TM2* (control) larvae were reared at 17°C under standard conditions until ecdysis to the third instar, and where then moved to 10% O_2_ at 24°C and left to pupate. We collected digital images of the pupal cases and measured the area of the pupal case when viewed from the dorsal aspect using ImageJ ^49^.

### TGPH Localization

*P{tGPH}* larvae were reared as described above until ecdysis to the third larval instar and either maintained at 21% O_2_ or transferred to 10% O_2_. After 12h, larvae were collected, and their fat bodies were dissected, fixed, stained and imaged using previously published protocols ^48^.

### Measurements of Growth

Females were left to oviposit on 50mm diameter Petri dishes contain standard cornmeal-molasses medium for 24h. Larvae were left to develop to the third larval instar and staged at the third larval instar by collecting newly ecdysed third instar larvae and transferring them to fresh food plates, in 3h cohorts. Larvae were then maintained at either 21% O_2_ or 10% O_2_. Larval mass was measured every 6h up to 24h after ecdysis to the third larval instar (AEL3) by removing thirty larvae from the food plates and massing them individually on a Mettler Toledo XPR Microbalance. Larvae were not returned to the food plate but were used to measure gene expression (see below).

For final body size, we collected and measured digital images of pupal cases from larvae reared throughout the third larval instar at 21% and 10% O_2_.

### Quantitative PCR (qPCR)

Larvae were collected every 6h for the first 24h of the third larval instar, as described above, and preserved in groups of ten in RNAlater (ThermoFischer Scientific). RNA was extracted using Trizol (Invitrogen, Grand Island, NY, USA) and treated with DNase I (ThermoFischer Scientific). RNA was quantified with a NanoDrop One (ThermoFischer Scientific) and reverse transcribed with High-Capacity cDNA Reverse Transcription Kit (Applied Biosystems, Carlsbad, CA, USA). Quantitative RT–PCR was conducted using PowerUp SYBR Green Master Mix (Applied Biosystems) and measured on an QuantStudio 3 Real-Time PCR system (Applied Biosystems). mRNA abundance was calculated on three biological replicates of ten larvae, using a standard curve and normalized against expression of ribosomal protein 49 (RP49). Standard curves were generated using seven serial dilutions of total RNA extracted from the isogenic control line (60,000): 5x 1^st^ instar larvae, 5x 2^nd^ instar larvae, 3^rd^ instar larvae (male), 5x pupae (male), 5x adult flies (male). Primer sequences used in the study were:

RP49 Forward: AAG AAG CGC ACC AAG CAC TTC ATC RP49

Reverse: TCT GTT GTC GAT ACC CTT GGG CTT

Imp-L2 Forward: GCG CGT CCG ATC GTC GCA TA

Imp-L2 Reverse: TTC GCG GTT TCT GGG CAC CC

4EBP Forward: CAG ATG CCC GAG GTG TAC T

4EBP Reverse: GAA AGC CCG CTC GTA GAT AA

E74B Forward: ATC GGC GGC CTA CAA GAA G

E74B Reverse: TCG ATT GCT TGA CAA TAG GAA TTT C

## Acknowledgements

This work was funded by the National Science Foundation [IOS 1122157 to J.F.H. and IOS 0845847 to A.W.S.]

## Author Contributions

GPK, VC, JFH and AWS conceived the study and designed the experiments. GPK, VC, HB, PT and SJLG conducted the experiments and analyzed the data. GPK, VC, JFH and AWS wrote the manuscript.

## Extended Data Figures

**Supplementary Figure 1:**
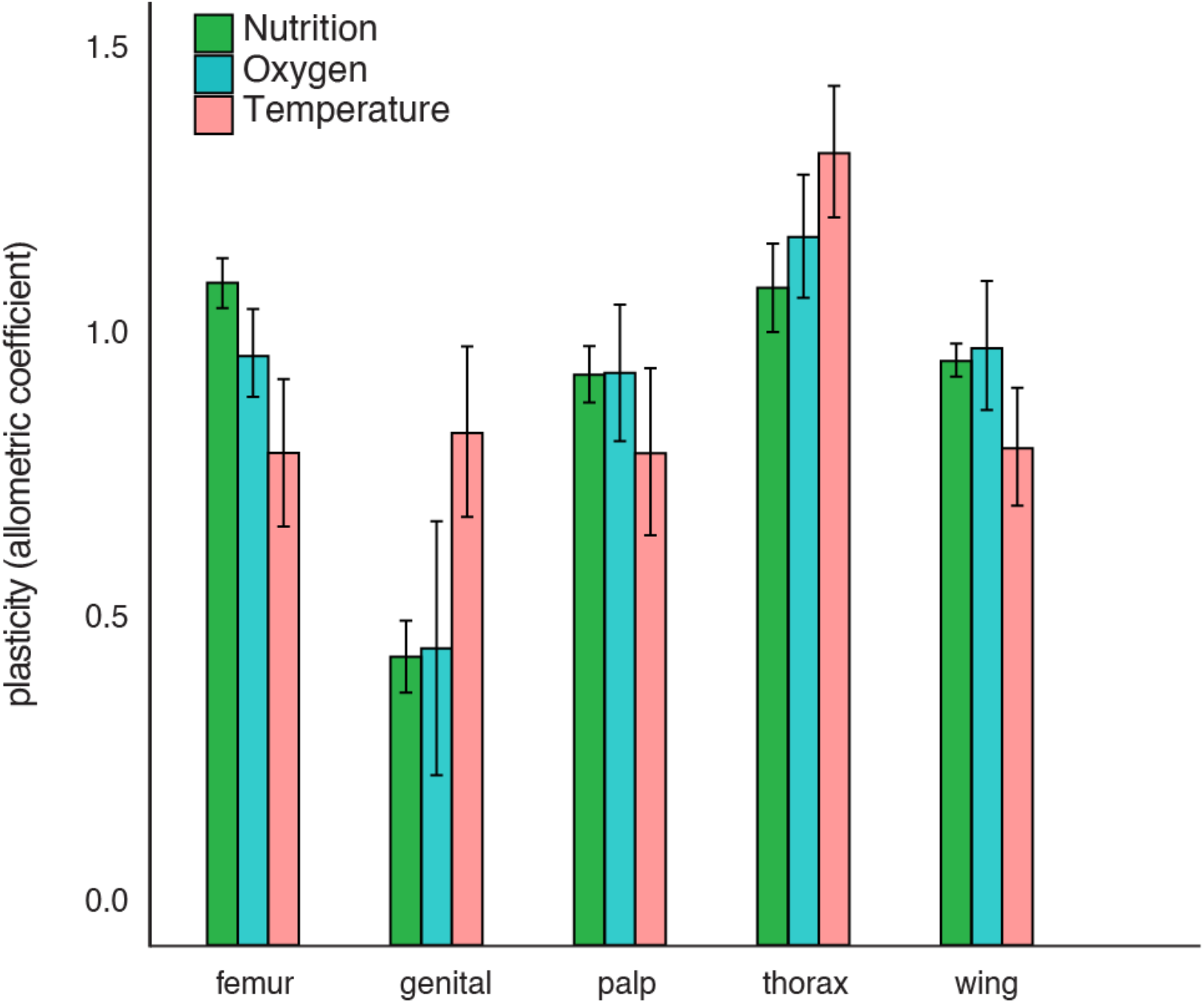
Relative plasticity of different traits to environmental variation in developmental nutrition, oxygen and temperature. The allometric coefficient captures the extent to which trait size varies relative to the body as a whole; that is, relative plasticity. An allometric coefficient of 1 indicates that the trait scales isometrically with overall body size. Error bars are 95% confidence intervals.

**Supplementary Figure 2:**
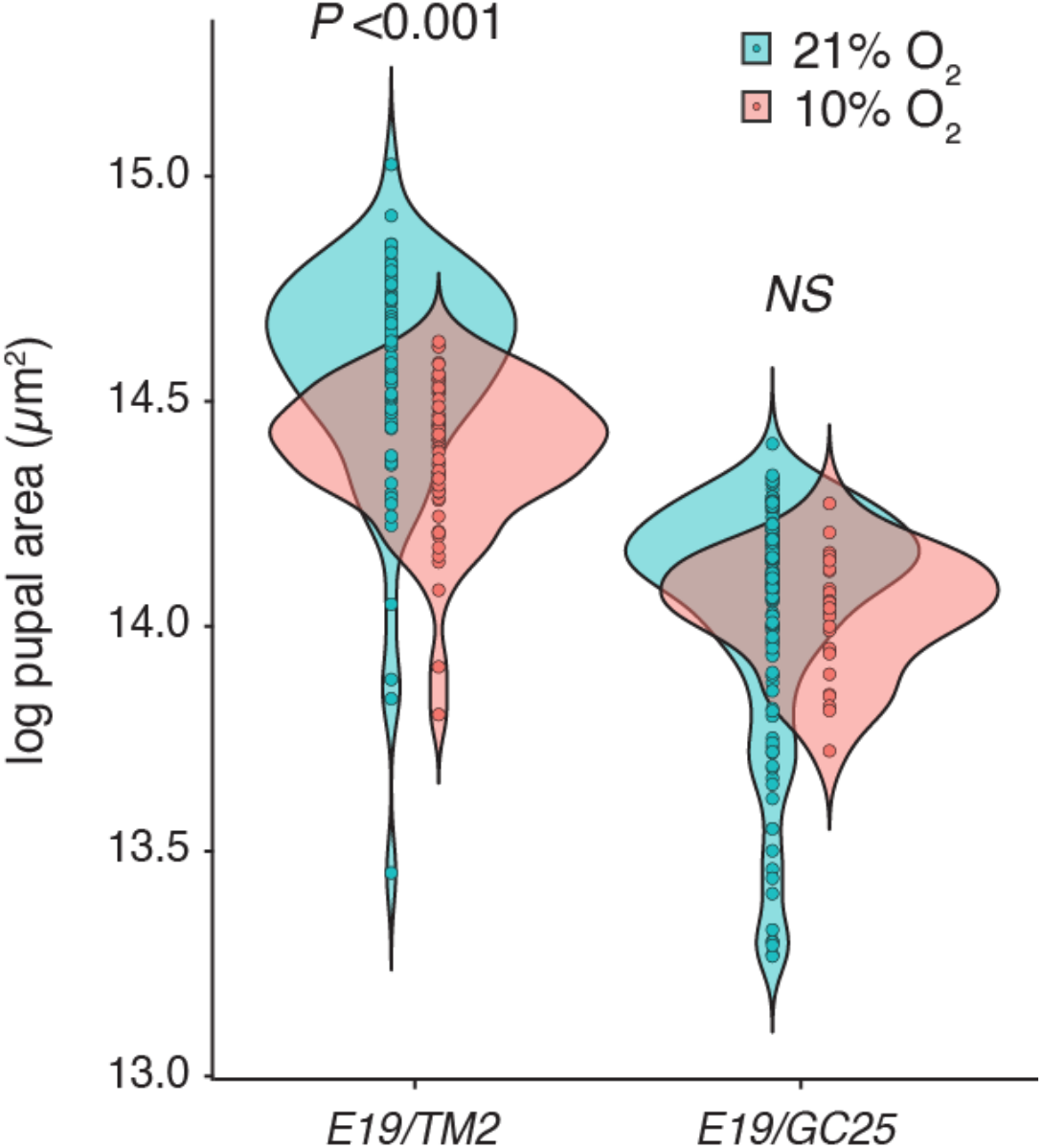
Suppression of the Insulin-Signaling pathway is necessary to inhibit growth under hypoxia. Adult body size shown in log pupal case area in flies functional IIS (*E19/TM2*) and flies with suppressed IIS (*E19/GC25*). Mean pupal size for each genotype across oxygen levels was compared using a pooled *t*-test. NS: not significant at *P*> 0.05.

**Supplementary Figure 3:**
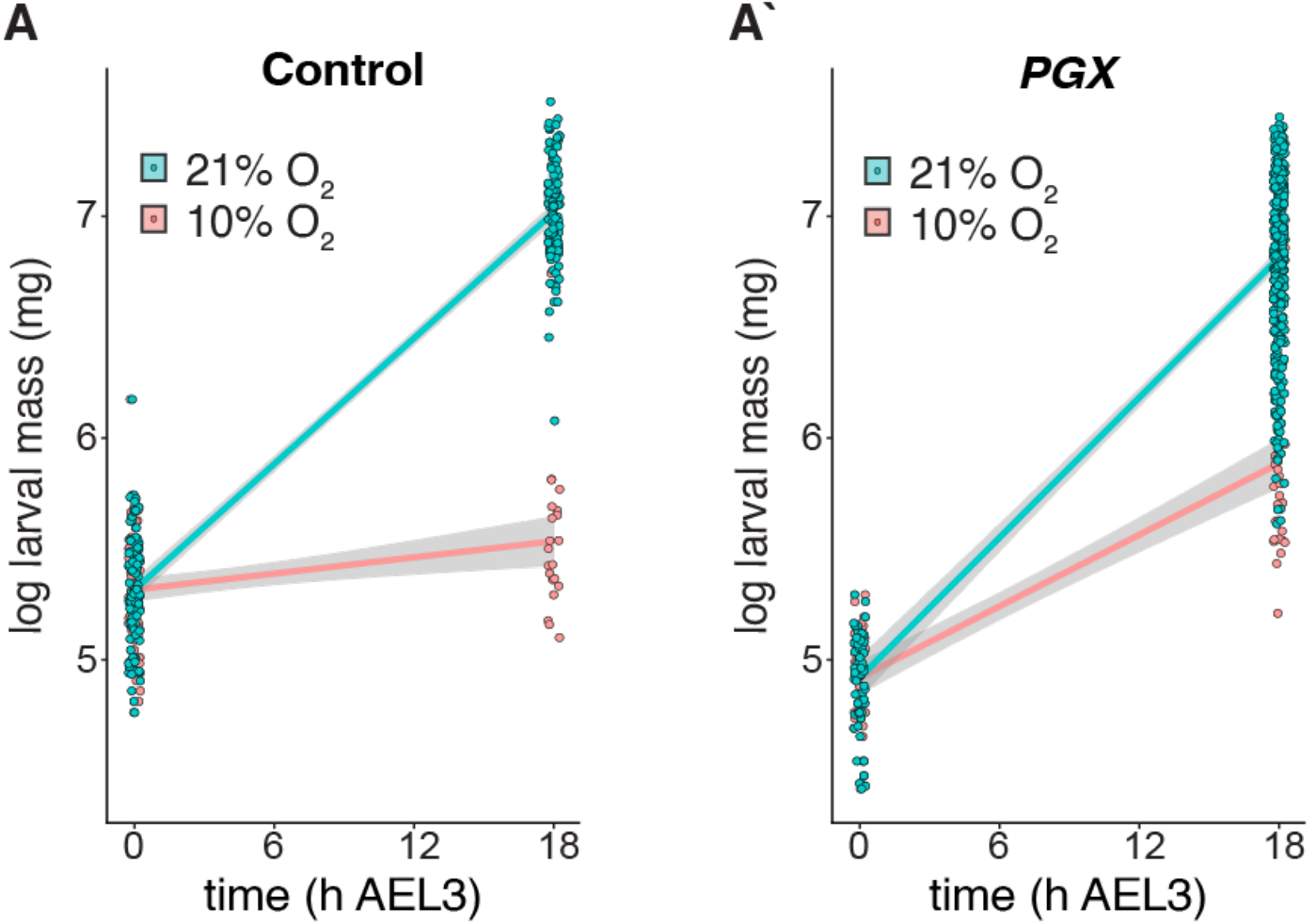
Ablation of the PG (site of ecdysone production) alleviates negative effects of hypoxia on growth. (A) 3^rd^ instar larval weight in log-mass over 18 hours in wildtype flies and (A’) *PGX = phm>grim;tubGAL80*^*TS*^ flies.

**Supplementary Figure 4:**
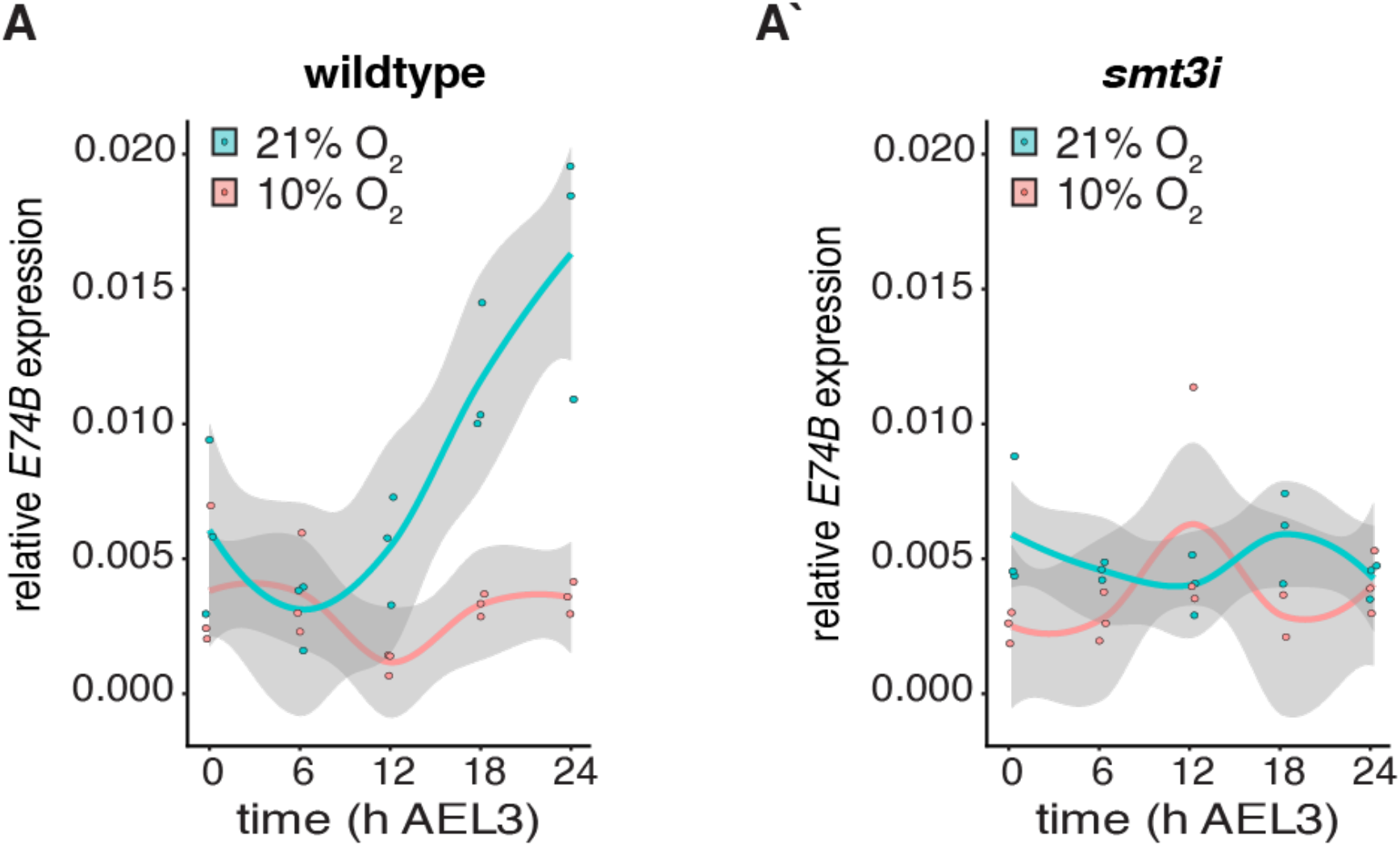
*E74B* mRNA expression. Transcript levels of *E74B* over 24 hours after L3 eclosion in control larvae were determined with qPCR, normalized to RP49 mRNA. (A’) Transcript levels of *E74B* in *smt3i* larvae. Grey bands are 95% confidence limits around the mean.

